# Physiological adaptations of leaf litter microbial communities to long-term drought

**DOI:** 10.1101/631077

**Authors:** Ashish A. Malik, Tami Swenson, Claudia Weihe, Eric Morrison, Jennifer B. H. Martiny, Eoin L. Brodie, Trent R. Northen, Steven D. Allison

## Abstract

Drought represents a significant stress to soil microorganisms and is known to reduce microbial activity and organic matter decomposition in Mediterranean ecosystems. However, we still lack a detailed understanding of the drought stress adaptations of microbial decomposers. We hypothesised that drought causes greater microbial allocation to stress tolerance relative to growth pathways. Here we present metatranscriptomic and metabolomic data on the physiological response of *in situ* microbial communities on plant leaf litter to long-term drought and pulse wetting in Californian grass and shrub ecosystems. Wetting litter after a long dry summer caused only subtle shifts in gene expression. On grass litter, communities from the decade-long ambient and reduced precipitation treatments had distinct functional profiles. The most discernable physiological adaptations to drought were production or uptake of compatible solutes to maintain cellular osmotic balance, and synthesis of capsular and extracellular polymeric substances as a mechanism to retain water. The results show a clear functional response to drought in grass litter communities with greater allocation to survival relative to growth that could affect decomposition under drought. In contrast, communities on chemically more diverse and complex shrub litter had smaller physiological differences in response to long-term drought but higher investment in resource acquisition traits across treatments, suggesting that the functional response to drought is constrained by substrate quality. Our findings suggest, for the first time in a field setting, a trade-off between microbial drought stress tolerance, resource acquisition and growth traits in leaf litter microbial communities.

## Introduction

Drought is common in terrestrial ecosystems, and climate change is making drought more frequent and severe^1,2^. Mediterranean ecosystems like those in California, USA, that experience summer drought are particularly vulnerable to climate change through reduced precipitation and increased evapotranspiration thus intensifying drought effects^3^. Drought affects soil processes through multiple mechanisms including limitations to resource diffusion and transport as well as organismal physiological responses to water stress^4–7^. Both are known to cause a decline in growth and activity of soil microorganisms^6,8^. As a direct effect of water limitation, microorganisms use their cellular machinery to maintain osmotic balance with the surrounding environment which involves intracellular accumulation of solutes or altering of the cell envelope to retain water^5,6,9,10^. Water limitation in soil also affects microbial growth and survival indirectly by altering substrate transport and cellular motility^6^. However, we do not have a thorough understanding of the key physiological adaptations of *in situ* microbes to drought thus demanding more mechanistic studies at individual population, collective community, and ecosystem levels. The knowledge gaps in our understanding of drought impacts introduces uncertainty in predictions of ecosystem processes under environmental change.

Summer drought in mediterranean ecosystems ends when seasonal precipitation begins in winter. The first rainfall after a long dry period stimulates microbial revival in plant litter and the underlaying soil^6,11,12^. The wet-up induced flushes of carbon and nutrients present a significant change in growth conditions for microorganisms^13–15^. Such drying-rewetting cycles cause fluctuations in water potential that are known to cause alterations in terrestrial nutrient pools and fluxes^6,13,16^. It has been demonstrated that the rapidly initiated CO_2_ pulse following the first rainfall after a long summer drought is linked to resuscitation and mortality of certain taxa of microorganisms^11,12,14^. However, the functional impacts of wet-up in terms of changes in gene expression, protein synthesis or metabolite production at the organismal and community level still remain unclear^17,18^.

Long-term drought selects for stress tolerant microorganisms but it also alters plant communities^19^ and therefore plant litter chemistry that in turn can select for different microbial communities in the litter layer and the soil below^7,12,20,21^. Selection based on the chemical quality of litter substrates could affect community physiology related to resource acquisition (substrate discovery, breakdown and uptake) and will likely impact drought tolerance and overall fitness of populations in the community^22^. Thus, the indirect effects of drought via changes in plant litter chemistry could significantly modify collective community physiology and therefore ecosystem process rates^7,21^. An assessment of drought impact on microbial physiology and its biogeochemical implications must therefore also consider these indirect effects.

The overall aim of this study was to identify the physiological mechanisms of drought stress response of microbial communities on plant leaf litter—the surface layer of soil. We investigated the stress physiology of distinct leaf litter communities arising from grassland and shrubland ecosystems undergoing field precipitation manipulation since 2007, with the drought treatment involving a ~40% reduction in annual precipitation that results in reduced decomposition rates^7^. Litter bags with litter from the different treatments (grass or shrub litter from ambient or reduced precipitation treatments) were deployed in the respective plots in summer 2017 and *in situ* wetting was performed at the end of the dry season to simulate commencement of seasonal precipitation (Figure 1). This approach was aimed at assessing microbial functional response to pulse wetting and subsequent drying, characteristic of these Mediterranean ecosystems.

**Figure 1:**
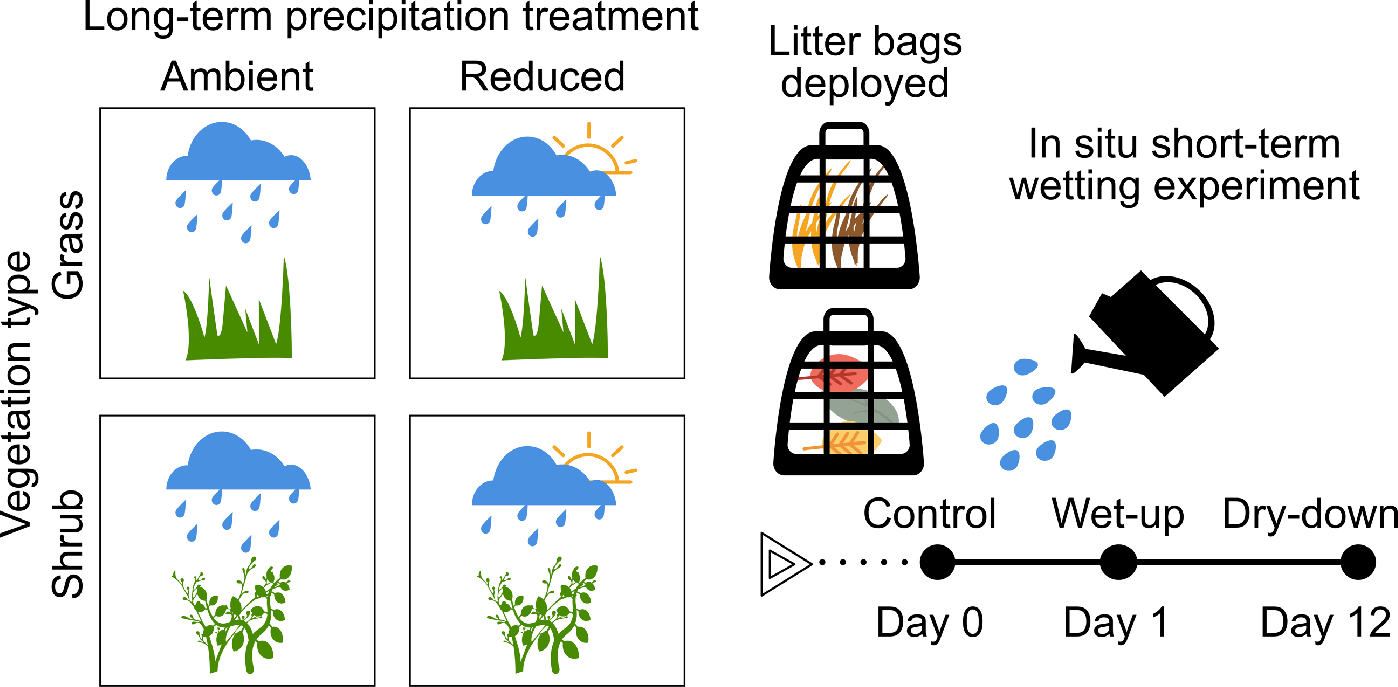
Experimental design and sampling scheme. *In situ* wetting was performed on litter bags deployed at the Loma Ridge Global Change Experiment that involved field scale precipitation manipulation (ambient and reduced) for a decade under two vegetation types (grass and shrub). Litter bags (n=4) were harvested before, one and 12 days after a discrete wetting event.

We hypothesised that long-term drought causes greater microbial allocation to stress tolerance relative to growth pathways. In the absence of chronic stressful conditions like drought, microbial communities should divert less investment into stress tolerance and therefore more to growth. More specifically we hypothesised that (1) long-term drought causes increased gene expression and metabolite production associated with osmoprotection and stress response, (2) the functional resuscitation responses to wetting are smaller in drought communities compared to ambient, and (3) chemically diverse and complex shrub litter requires increased investment in resource acquisition pathways which constrains allocation to drought stress response. We analysed the microbial physiological response using metatranscriptomics (community gene expression) and metabolomics (endo and exometabolites) to analyse community acclimation to acute changes in water potential as well as the adaptations to chronic drought stress. Our results demonstrate multiple physiological adaptations within drought communities that enable survival in such environments.

## Results

### Litter microbial community respiration on wet-up and dry-down

In this study, community-aggregated traits (means of functional traits found in a given community^23^) were used as a proxy to assess collective community physiology and its potential implications for ecosystem functioning. Community-level functional impacts of *in situ* pulse wetting were assessed in terms of changes in rate of respired CO_2_, gene expression and metabolite production. Pulse wet-up led to an increase in CO_2_ respiration which returned to the initial rate after 12 days of dry-down (Supplementary Information, Figure S1). There was no discernable difference in this dynamic between vegetation and precipitation treatments (two-way factorial ANOVA; p>0.05). Local conditions like canopy cover over litter bags and shading were probably more important than litter type or historical precipitation levels in determining the variation in moisture levels in litter after 1 day. Respiration rate correlated strongly with litter moisture content (Figure S1).

### Community functional and taxonomic diversity

Multivariate ordination analysis of transcripts (at the level of function in Subsystems classification) and metabolites suggested that wet-up and dry-down treatments did not change the functional composition of communities (Figure 2a-b; PERMANOVA p>0.05). Results across these treatments indicated that grass and shrub communities were functionally dissimilar (Figure 2a-b). Shrub litter communities demonstrated higher transcript functional α diversity than grass communities (Figure 2c). Effects of long-term reduced precipitation on microbial functioning were variable in the two vegetation types. On shrub litter, communities from reduced and ambient precipitation treatments shared more transcripts and metabolites than in grass communities where reduced precipitation clearly altered microbial physiology (Figure 2a-b). Transcript functional α diversity was similar in shrub communities from ambient and reduced precipitation treatments (Figure 2c). However, we observed a higher functional diversity of transcripts in grass communities from reduced precipitation treatment in comparison to ambient (Figure 2c).

**Figure 2:**
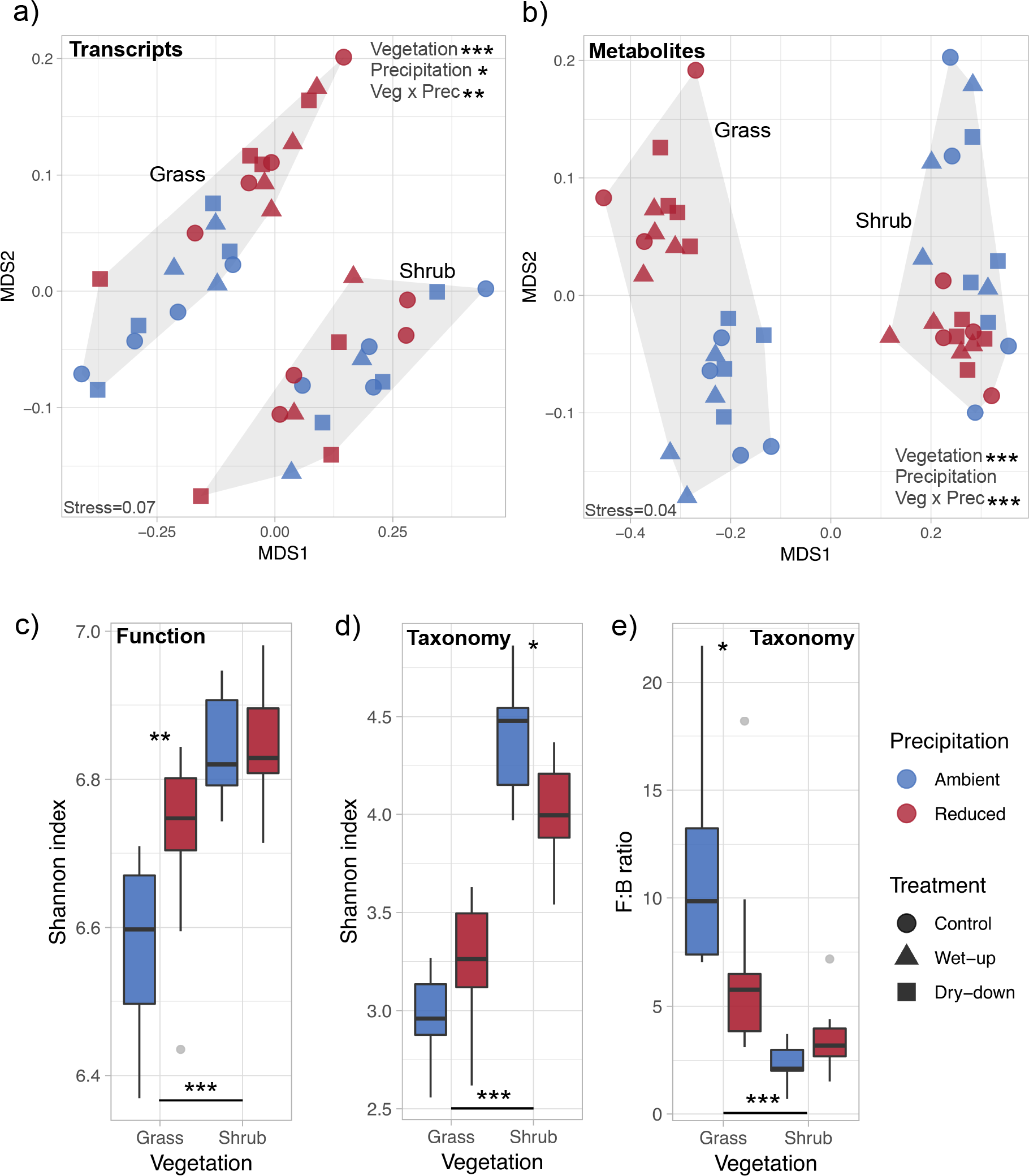
Community functional and taxonomic diversity. Two dimensional NMDS ordination of (a) transcripts (function) and (b) metabolites. Treatments from the *in situ* wetting experiment involved control (day 0), wet-up (day 1) and dry-down (day 12). α diversity of (c) functions and (d) taxa derived from transcripts across vegetation and long-term precipitation treatments; n=9-12. (e) Community fungal:bacterial (F:B) ratio estimated as the ratio of mRNA sequences assigned to fungi and bacteria; n=9-12. In a-b asterisks mark the significance of treatments that cause clustering of similar samples based on Bray–Curtis dissimilarity index analysed using permutational multivariate analysis of variance (PERMANOVA), and in c-e asterisks mark the significance of differences between groups as analysed by Tukey’s multiple comparison test (*** p < 0.001, ** p < 0.01, * p < 0.05). No statistical tests on impacts of short-term wet-up and dry-down treatments were significant.

We used the taxonomic annotations of functional genes obtained from metatranscriptomics to ascertain the taxonomic α diversity and composition of the active litter microbial communities that consisted of bacteria and fungi. Communities from shrub vegetation had significantly higher taxonomic α diversity of transcripts than grass communities (Figure 2d). Within each vegetation type, communities from ambient and reduced precipitation treatments differed taxonomically (Figure S2), demonstrating a clear effect of simulated long-term drought in shaping the taxonomic composition of the microbial community. Short-term wet-up and dry-down treatments did not change the taxonomic composition of communities (Figure S2). Fungi dominated the active communities in both grass and shrub vegetation. Fungal:bacterial ratios obtained from transcript annotations as well as from ribosomal RNA concentrations (Bioanalyzer-derived 18S:16S rRNA ratio) were higher in grass communities (Figure 2e, S3, S4). Relative abundance of fungi in communities was higher in grass litter with a history of ambient precipitation compared to reduced precipitation (Figure 2e). In shrub communities, there was no discernible shift in fungal:bacterial ratio in response to long-term precipitation treatment (Figure 2e).

### Impact of long-term drought on gene expression

We aimed to uncover the physiological mechanisms that underlie microbial adaptations to long-term drought by analysing variable patterns in gene expression across the decade-long simulated precipitation treatments. The overall proportions of functions identified from metatranscriptomics were largely similar in communities from the two litter types (Figure 3a). In grass communities, reduced precipitation caused systematic shifts in many functional classes (Figure 3b). Functional classes that were significantly higher in relative abundance under reduced precipitation treatment were membrane transport, iron acquisition and metabolism, phosphorus metabolism, stress response, DNA metabolism, and cell division and cell cycle. Classes with significantly lower relative abundance under reduced precipitation treatment compared to ambient were secondary metabolism, respiration and nitrogen metabolism. These shifts in functional classes imply higher expression of conventional housekeeping genes in grassland ambient than reduced precipitation treatment. However, in shrub communities there were only small, non-significant changes in abundances of these functional classes in response to the 10-year reduced precipitation treatment (Figure 3b).

**Figure 3:**
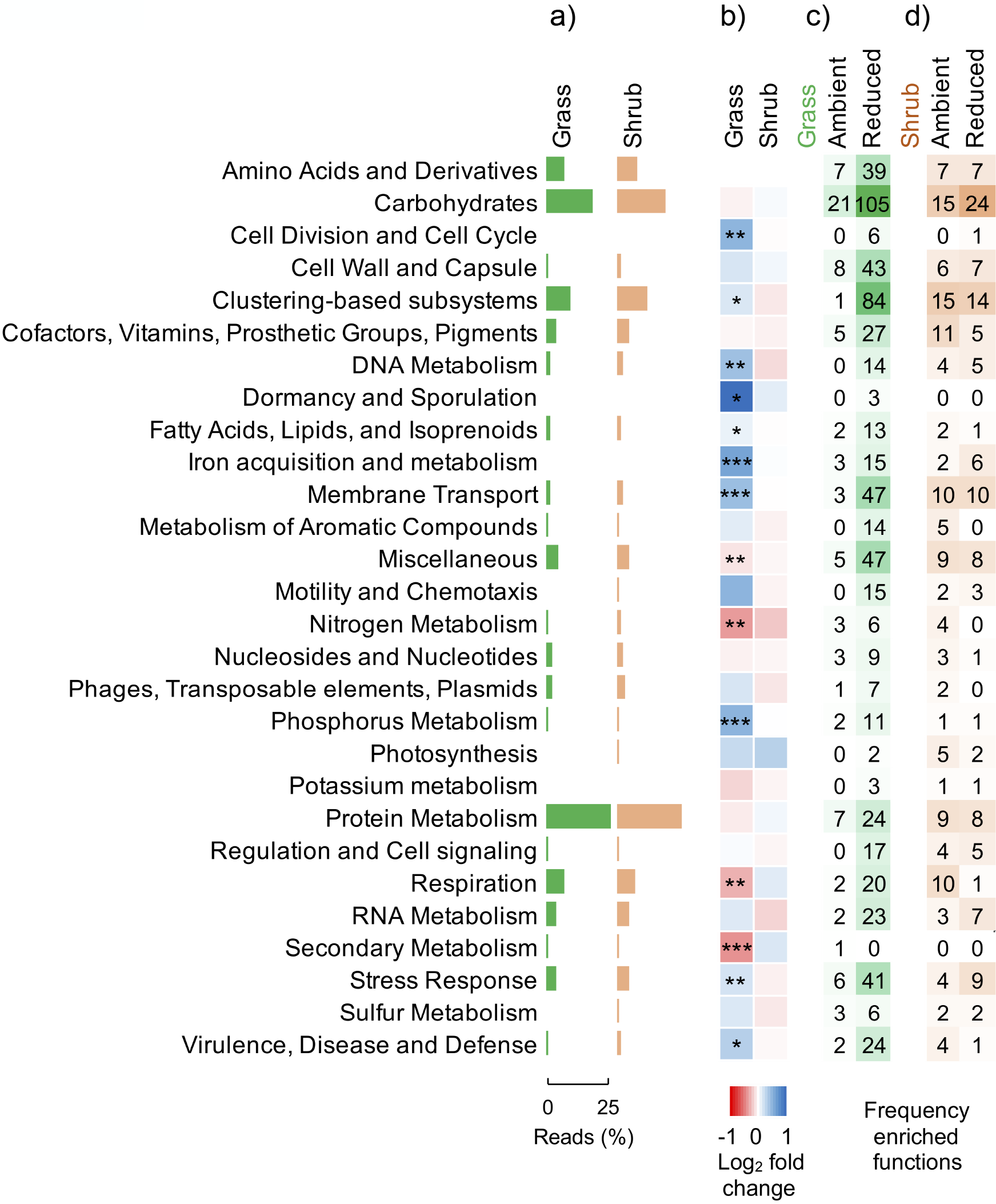
Impact of long-term drought on microbial physiology. (a) Mean relative abundance of transcripts at level 1 of Subsystems classification in grass and shrub litter communities (n=18-24). (b) Fold change in gene expression (at level 1 class) in communities from reduced precipitation relative to ambient control. Asterisks mark significant drought-induced shifts in functional groups as analysed by one-way ANOVA (*** p < 0.001, ** p < 0.01, * p < 0.05; n=9-12). Frequency of significant transcript indicators (p<0.05) in the upper level functional groups (level 1 class) across precipitation treatment in (c) grass and (d) shrub communities (n=9-12). These indicators were unique to or enriched in communities in either ambient or reduced precipitation treatments and represent functional indicators of either treatments.

Drawing conclusions from shifts in upper-level classes of functions may be misleading as this level consists of a myriad genes whose expression may decrease or increase in response to the environmental perturbation^24^. Multiple genes belonging to the same class responding in opposite directions may cancel out resulting in no net effect for that class of functions. Therefore, we also assessed shifts in expression of individual genes to identify those that were unique to or significantly enriched in communities from either ambient or reduced precipitation treatments. From the 7140 Subsystems functions that were annotated in our dataset, in grassland communities we identified 665 transcript indicators of reduced precipitation treatment but only 87 indicators of ambient treatment (Figure 3c, Data S1). In shrub litter communities, we observed similar numbers of indicator transcripts in contrasting precipitation treatments, with 140 under ambient precipitation and 120 under reduced precipitation (Figure 3d, Data S1).

The majority of transcript indicators of reduced precipitation treatment in grass litter communities could be directly or indirectly linked to physiological adaptations to moisture stress. In contrast, most of the indicators of ambient treatment (Figure 3c) belonged to common housekeeping functions (level 1 class carbohydrates: 21 functions). The highest number of transcript indicators of reduced precipitation in grass litter also belonged to the level 1 class of carbohydrates (Figure 3c, 105 functions) but represented functions linked to metabolism of mono-, di- and oligosaccharides such as L-rhamnose, trehalose, maltose and maltodextrin, D-galacturonate, etc., and organic acids such as malonate. While some of these can be linked to conventional or alternative housekeeping functions, increased expression of genes linked to metabolism of sugars like trehalose suggests an adaptive mechanism of compatible solute accumulation to maintain cellular osmotic balance^5,10,25,26^.

A significant number of indicators of reduced precipitation in grass litter also belonged to the classes of membrane transport (47 functions) or stress response (41 functions) which were almost absent in the indicator profiles of ambient communities (Figure 3c). Membrane transport functions were mostly annotated to multi-subunit cation antiporters (Na+ H+ antiporters), Ton and Tol transport systems, ABC transporters and protein secretion systems. Increased expression of genes for cation antiporters indicates a drought acclimation strategy of accumulating inorganic ions aimed at maintenance of cellular osmolarity^6,9,27^. In the level 1 class of stress response, we observed increased expression of genes related to uptake/biosynthesis of choline, betaine and ectoine. These metabolites are widely-reported osmolytes in plants and microorganisms^6,9,10,25,28^ and provide additional evidence for osmolyte accumulation as an adaptation mechanism in drought communities.

We also observed enrichment of transcripts belonging to the cell wall and capsule class in communities from reduced precipitation in grass litter (43 functions, Figure 3c). Significant gene indicators here were linked to metabolism of capsular and extracellular polysaccharides (EPS), Gram-negative cell wall components (lipopolysaccharide assembly), and peptidoglycan biosynthesis demonstrating strategies either to retain water through capsular or EPS “sponges”, or to lower the permeability of cell walls to avoid water loss^6,9,26^. Genes of the classes RNA and protein metabolism that were significantly higher in reduced precipitation treatments were related to functions like transcription, RNA processing, and protein synthesis/modification and were mostly of bacterial origin, whereas similar transcript indicators in ambient communities were mostly eukaryotic. This evidence links these functional indicators to drought-induced shifts in fungal:bacterial ratios of microbial communities (Figure 2e)^7^. A high number of functional indicators belonged to Clustering-based Subsystems (84 functions, Figure 3c) and miscellaneous (47 functions), but the majority of these were unassigned or putative functions that did not provide any clear functional insight.

Shrub litter communities were functionally more diverse than grass litter communities but precipitation treatment did not change the functional diversity (Figure 2a-c). Correspondingly, we observed a similar number of unique functions across the contrasting precipitation treatments within shrub litter (Figure 3d). Unlike grass communities, very few distinctive stress response functions were identified in shrub communities under reduced precipitation treatment; these were linked to trehalose metabolism, oxidative stress and multi-subunit cation (Na+ H+) antiporters. Thus, the drought stress response physiology of microbial communities is variable and could depend on other traits like resource acquisition linked to litter chemistry.

### Long-term drought impact on community metabolite production

We next identified metabolites that were unique to or enriched in either reduced or ambient precipitation treatments. The results corroborated the patterns revealed using metatranscriptomics (Figure 4, S5). Ectoine, used as a compatible solute to maintain internal water potential, showed greater abundance in the reduced precipitation treatment particularly in grass litter communities. 5-oxo-proline, also used as an osmolyte, showed greater abundance in shrub litter communities under reduced precipitation treatment. On the contrary, in the grass litter under ambient precipitation treatment, various biogenic amino acids such as aspartic acid, and purine ribonucleosides such as adenosine and guanosine were enriched; these are molecular building-blocks of proteins and RNA, respectively and indicate growth and biosynthesis. These differential metabolite profiles in contrasting precipitation treatments in grass communities corroborate the metabolic tradeoff between growth and water stress adaptations suggested by the transcriptomics data.

**Figure 4:**
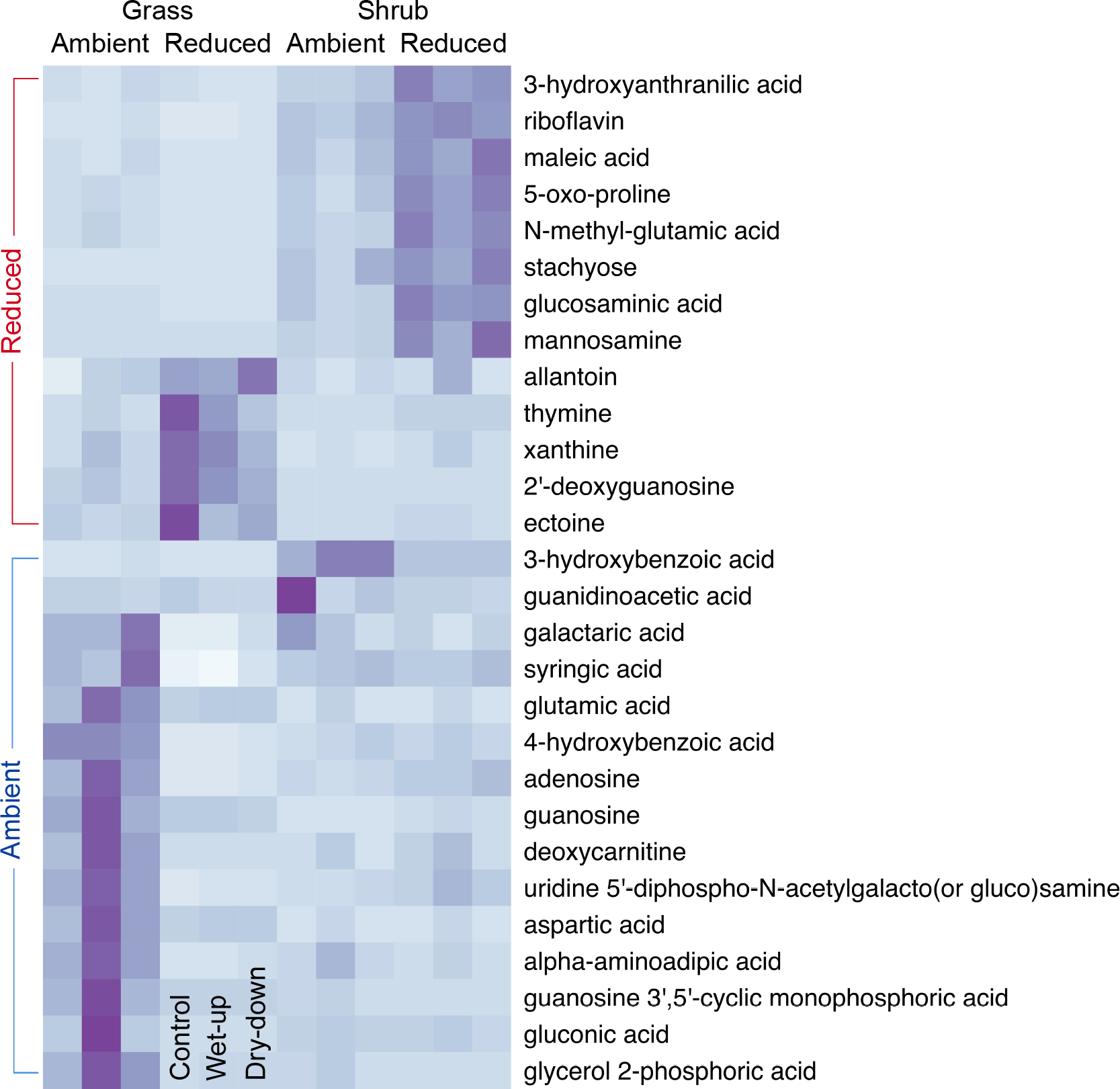
Metabolite abundance across vegetation and precipitation treatments. Heatmap showing mean peak heights (n=3-4) which relates to the abundance of metabolites that were significantly higher (p<0.05) in either ambient or reduced precipitation treatments across both litter types. Rows of metabolites are clustered horizontally according to vegetation and precipitation treatments.

Consistent with metabolic tradeoffs, we observed 47 and 27 significant negative correlations between metabolite pairs (out of 378 tested) in grass and shrub litter communities, respectively (Figure S6). In grass litter communities, we observed significant negative correlations between key growth indicators aspartic acid and adenosine and drought stress indicator ectoine (Figure 5). In shrub litter communities, although we did detect slightly higher amount of osmolytes ectoine and 5-oxo-proline in the reduced precipitation treatment, there were no negative correlations between these drought stress indicators and the growth indicators (Figure 5). This analysis demonstrates metabolic trade-offs between relevant growth and stress indicators in grass litter communities but not in shrub litter communities.

**Figure 5:**
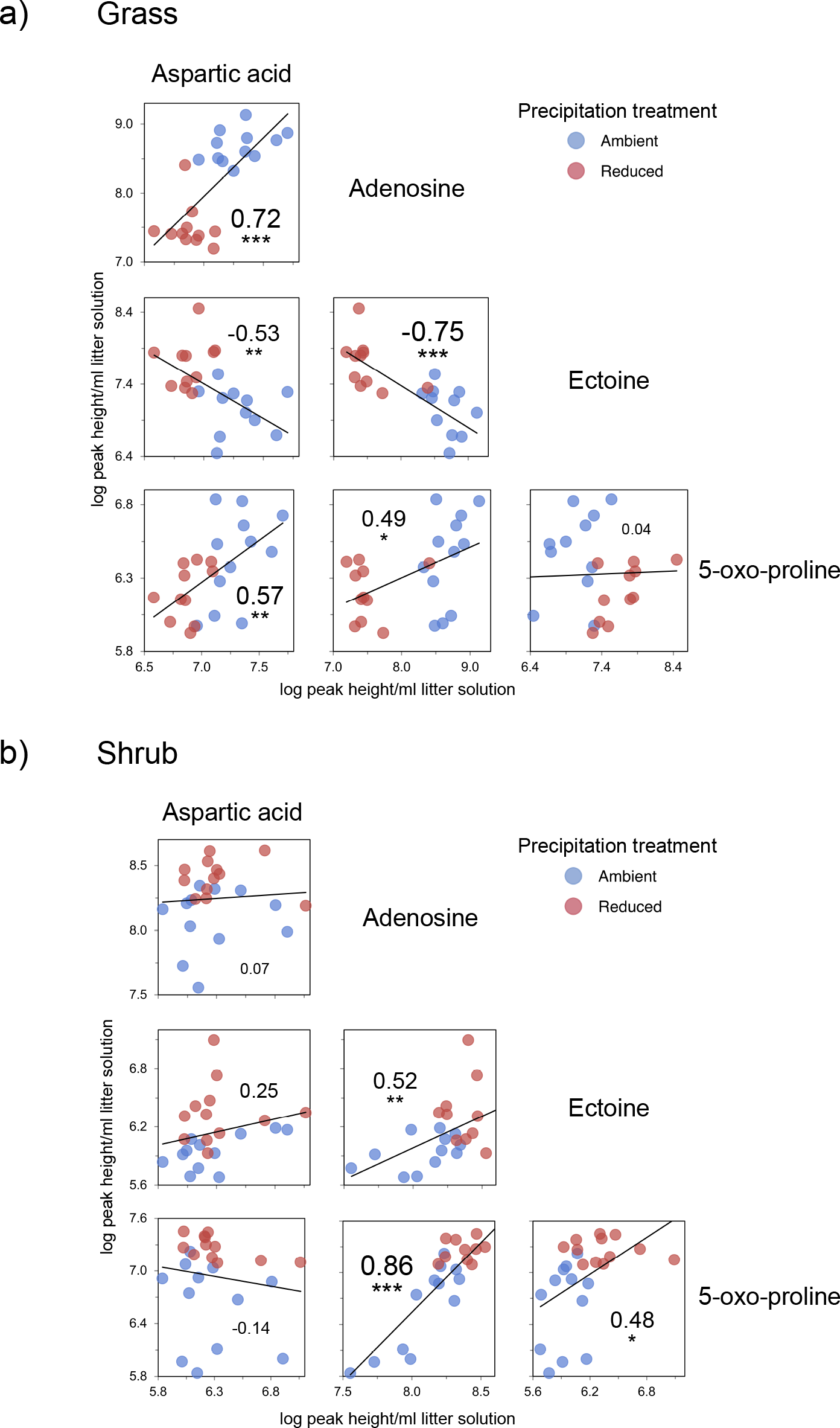
Correlations between metabolite abundances demonstrating metabolic tradeoffs. Linear regression trends of aspartic acid and adenosine (growth indicators), and ectoine and 5-oxo-proline (osmolytes as stress indicators) in (a) grass and (b) shrub litter communities. These metabolites were the most relevant from among those that were significantly enriched in either ambient or reduced precipitation treatments (shown in figure 4). Numbers within each scatter plot are correlation coefficients (r) and asterisks denote the significance of the relationship across treatments (*** p < 0.001, ** p < 0.01, * p < 0.05; n=23-24).

### Community functions unique to litter types

When shrub litter communities were compared to grass litter communities under ambient precipitation treatment, a large number of unique functions were observed in shrub communities (Figure S7, Data S2). Of the 182 functional indicators of grass communities, most belonged to the class protein metabolism (39 functions) and were linked to small and large subunit ribosomal protein synthesis as well as proteasome-mediated degradation of unneeded or damaged proteins^29^. On the contrary, we identified 1588 unique functions in shrub communities (Figure S7). The majority of functions unique to or enriched in shrub litter communities belonged to the class of carbohydrates (208 functions) and ranged from central carbohydrate metabolism to metabolism of mono-, di- and oligosaccharides, organic acids and sugar alcohols. In addition, 115 indicators of shrub litter communities were annotated to amino acid metabolism. The presence of a large number of functional indicators for carbohydrate and amino acid metabolism indicates increased investment in substrate degradation, uptake and assimilation in shrub communities. Microbial communities in shrub litter may have increased investment in these resource acquisition traits to degrade the more chemically complex and diverse substrates.

### Functional response to *in situ* wet-up

While there were no significant shifts in overall metatranscriptomics-derived functional as well as taxonomic diversity and composition in response to wetting and subsequent drying (Figure 2a-b, S8), we did observe changes in gene expression and metabolite production linked to certain functions. However, the functional response to wetting was similar in grass communities from ambient and reduced precipitation treatments (Figure 6a). In grass litter communities, genes for membrane proteins (ompA); amino acid, oligopeptide, and phosphate transporters (livG, livK/J, pstS, oppC); and flagellar motility (fla, flaA) were upregulated in response to wetting while those coding for biomolecular repair (rad51, hscA, hscB) were downregulated (Figure 6a-b). Outer membrane protein A (ompA) acts as a porin to allow slow passive penetration of small solutes^30^. Upregulation of genes for branched-chain amino acid ABC transporters (livG, livK/J) and the oligopeptide transport system (oppC; Figure 6a-b) suggests transporter-mediated excretion of compatible solutes to reduce intracellular solute concentrations in order to prevent cell lysis as external water potential rises^6,9,17,31,32^. Further evidence comes from the significant increase in abundance of amino acids including branched-chain amino acids like leucine and valine in the metabolite profiles on wetting (Figure S9). These patterns demonstrate specific physiological acclimation mechanisms that microbes express in response to wetting.

**Figure 6:**
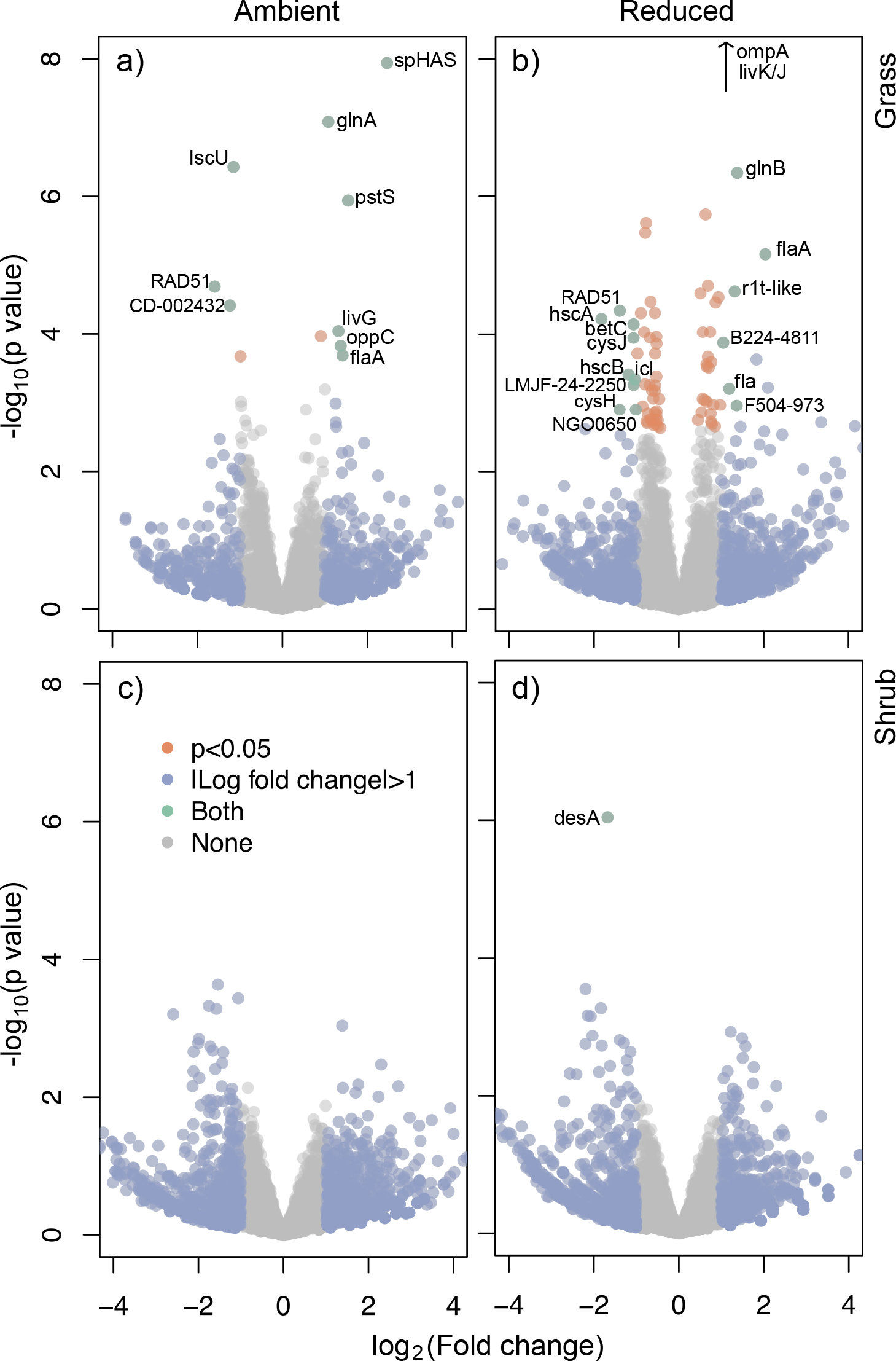
Gene expression patterns in response to *in situ* wetting. Volcano plots of control (day 0) vs. wet-up (day 1) litter showing transcript level changes in (a) grass ambient, (b) grass reduced, (c) shrub ambient and (d) shrub reduced precipitation treatments. Only genes with significant shifts in expression on wet-up with fold change more than double or less than half the control samples were labelled. Relevant upregulated genes- ompA: outer membrane protein A; livG, livK/J: branched-chain amino acid ABC transporter; pstS: phosphate ABC transporter; flaA, fla: flagellin protein; oppC: oligopeptide transport system. Relevant downregulated genes- IscU: iron-sulfur cluster assembly enzyme; rad51: DNA repair protein; hscA, hscB: chaperone protein.

Increased expression of flagellin protein (fla, flaA) on pulse wetting suggests upregulation of functions linked to motility in the presence of water to move and explore resources. This process appears to be repressed in dry conditions as liquid water becomes scarce and metabolic activity shifts towards energy preservation^9,33^. DNA repair protein (rad51) essential for repairing damaged DNA as well as for protection of intact DNA was downregulated on wetting. Higher expression of genes related to these repair proteins under dry conditions represents a stress tolerance and alleviation mechanism. Also downregulated on wetting were chaperones (hscA, hscB) involved in maturation of proteins like iron-sulphur cluster assembly (iscU). Stress-induced disruption of proteins impairs major biosynthetic and transport pathways and chaperones aid in mitigation of protein damage^9,30,34^.

No gene groups with significant changes in expression patterns with wetting were identified in shrub communities (Figure 6c-d). Metabolite profiling also confirmed that wetting and drying had less impact on physiological processes in shrub compared to grass litter communities (Figure 2b, S5, S9). However, abundance of biogenic amino acids such as tyrosine, histidine and lysine as well as nucleotide bases such as guanine and cytidine were significantly higher on wetting in both grass and shrub communities (Figure S9). Elevated amounts of these amino acids and nucleotide bases that are precursors for protein and nucleic acid synthesis suggest increased microbial growth or turnover with higher water availability.

## Discussion

We demonstrated distinctive physiological adaptations of microbial communities growing on plant leaf litter in response to simulated decade-long drought. The differing gene expression and metabolite patterns across long-term precipitation treatments could reflect the differing genetic potential of the microbial communities, a consequence of drought-induced changes in microbial community composition^7,35,36^. The observations of community shifts suggests environmental pressures that select for taxa that can survive the physiological stress imposed by drought^36–39^. The community shifts could also be ascribed to the indirect effect of drought-induced changes in plant litter chemistry acting as a selection pressure^7,21^.

For the first time in a field setting, our data demonstrate up-regulation of stress-response genes and production of related metabolites in leaf litter microbial communities in response to drought. The most discernible physiological adaptation to drought was intracellular accumulation of organic osmolytes aimed at equilibrating the internal osmotic potential with that of its surroundings. We found some evidence to suggest that the flow of inorganic ions across the membrane was also regulated to maintain osmolarity. Gene expression analysis of *in situ* communities suggests that various compatible solutes like ectoine, betaine, choline and trehalose were employed as osmolytes^6,10,25^. Corroborating this inference, under reduced precipitation, we observed significantly higher concentrations of ectoine in grass litter communities and higher concentrations of choline and betaine in shrub litter communities. Another significant physiological adaptation was synthesis of capsule and EPS that enable cells to form biofilms and retain water for longer periods in drier environments^6,9,40^.

In ambient precipitation treatments, we observed more growth-related indicators which suggests that under less stressful conditions, microbial communities divert fewer resources into stress tolerance and therefore grow better^5,41^. Our data demonstrate that physiological response to drought trades off against growth by diverting investment away from central metabolism and associated assimilatory pathways such as amino acid and nucleotide synthesis. Physiological adaptation mechanisms like osmolyte accumulation can be energetically very expensive—by some accounts osmotic stress has been estimated to reduce growth yields by nearly 90%^5,42^. Such massive shifts in resource allocation can have big impacts on ecosystem C and N balance. To this end, we can relate increased stress investment in grass litter communities to decreased microbial decomposition under drought previously observed in the same field experiment^7^. However, stress response and growth indicators were largely similar in shrub ambient and reduced precipitation treatments.

We also revealed communities’ short-term acclimation strategies in response to pulse wetting and subsequent drying. However, there was no evidence that the functional resuscitation response to wetting was smaller in communities from long-term drought compared to those from ambient conditions. Wetting leaf litter after a long dry summer caused only subtle shifts in gene expression. It is possible that a single discrete wetting event is not sufficient to elicit a large shift in gene expression and overall physiology^11,36^. There could be high cellular costs of such physiological changes and cellular systems may not be primed to respond until sustained wet conditions appear.

Here we reported community-aggregated traits to understand the functional attributes of complex microbial communities in a precipitation manipulation. Although we observed a tradeoff between drought stress tolerance and growth traits in grass communities, shrub litter chemistry appears to alter these community-level tradeoffs. It is likely that the greater chemical complexity and recalcitrance of shrub litter (higher C:N ratio, higher proportion of lignin and other recalcitrant compounds and lower proportion of cellulose, hemicellulose and cell solubles than the grass litter^43^) makes it harder for microorganisms to degrade and imposes additional constraints on microbial physiology that affect stress tolerance strategies under drought. The chemically more diverse and complex shrub leaf litter supports a taxonomically and functionally more diverse decomposer community. We also observed a high frequency and diversity of functional indicators of substrate degradation, uptake, and assimilation in shrub litter communities that may reflect the low resource quality. This high level of investment in resource acquisition traits in shrub litter communities may have affected their response to drought stress if there are tradeoffs between resource acquisition and stress tolerance traits. Although such tradeoffs require further validation, we provide a framework to link ecosystem functioning with omics-derived traits of *in situ* microbial communities.

## Conclusions

Our results indicate that the metabolic costs of microbial physiological adaptations to drought trade off against growth traits. However, litter of poor chemical quality constrains gene expression and metabolite production associated with drought stress tolerance. Substrate quality and investment in resource acquisition traits can thereby alter the stress tolerance-growth trait relationship in microbial communities. The potential linkages of microbial traits with litter decomposition rates suggest that community-level trait trade-offs have consequences for organic matter decomposition, a key ecosystem process. Our data clearly identifies some of the physiological mechanisms of community-level adaptations to drought and possible trade-offs in stress tolerance, resource acquisition and growth. The significance of these trait trade-offs can be further explored by their representation in models of organic matter decomposition.

## Methods

### Field site description

The study site was part of the Loma Ridge Global Change Experiment situated near Irvine, California, USA (33°44’N, 117°42’E, 365 m elevation). The climate is mediterranean with mean annual temperature of 17°C and mean annual precipitation of 325 mm. Most precipitation falls between November and April, and a summer drought lasts from May to October. The vegetation at the site includes annual grassland adjacent to coastal sage shrubland which were the two litter types used in the experiment^7,19^. The grassland plots (6.7 × 9.3 m) consisted of exotic annual grasses such as *Avena*, *Bromus*, and *Lolium* and forbs such as *Erodium*, whereas the shrub plots (18.3 × 12.2 m) consisted of crown-sprouting shrub species such as *Salvia mellifera*, *Artemisia californica* and *Malosma laurina*. This shrub litter is known to have higher C:N ratio, higher proportion of lignin and other recalcitrant compounds and lower proportion of cellulose, hemicellulose and cell solubles than the grass litter^43^. For our experiment, we used a subset of plots with field precipitation manipulations that have been established since February 2007. Reduced precipitation treatment involved ~40% reduction in precipitation compared to ambient which was achieved by covering plots with clear polyethylene during a subset of rain events each winter. The design was replicated to have four plots per precipitation treatment (grassland ambient and reduced; shrubland ambient and reduced).

### Experimental design and sampling

Dry litter collected on 30 August 2017 from all four replicated plots within a treatment was pooled and homogenized. Litter bags were made by placing 6 g dry litter mass into 15 cm × 15 cm bags made from 1 mm mesh window screen. Bags were deployed on 12 September 2017. It was a particularly dry season and no sizeable precipitation was reported until December 2017. *In situ* pulse wetting was performed on 30 November 2017 to simulate a discrete rainstorm of ~13 mm using previously collected rainwater (Figure 1). Litter bags were collected before wetting to serve as dry control. Another batch of bags was collected the following day when most bags were still visibly wet. There was no precipitation in the following days and therefore the litter dried out. We collected another batch 12 days after wetting to study the dry-down response of microbial communities. 16 litterbags were sampled at each time point (2 vegetation types 2 precipitation treatments × 4 replicates). Retrieved litter bags were immediately transported to the lab at room temperature. A subsample of leaf litter was ground in a mixer (a quick whirl for 5 s) and used for RNA and metabolite extraction. 1g of intact leaf litter was placed in a 40ml glass vial and incubated for 1-2 h in the dark at room temperature (21°C) without manipulating moisture levels. Respired CO_2_ collected in the headspace was measured using EGM-4 Environmental Gas Analyzer for CO_2_ (PP Systems, Amesbury, MA, USA). Another subsample of intact litter was dried to constant mass at 65°C to obtain dry mass.

### RNA extraction and sequencing

We carried out RNA extraction on a litter aliquot of 0.2 g for shrub and 0.5 g for grass using RNeasy PowerSoil Total RNA Kit following manufacturer instructions (Qiagen, Hilden, Germany). Due to a high amount of organic compounds co-extracted from shrub litter, a lower amount of starting material was used in the extraction protocol to increase the RNA yield. After resuspending the RNA pellet in solution SR7 in the final step, purity and concentration of total RNA was assessed using a Bioanalyzer 2100 (Agilent, Santa Clara, CA, USA), Qubit fluorometer (LifeTechnologies, Carlsbad, CA, USA) and Nanodrop 2000 Spectrophotometer (Thermo Scientific, USA). Bioanalyzer-derived abundances of 18S and 16S rRNA in total RNA extracts was used to calculate fungal:bacterial (F:B) ratios. 200-500 ng of total RNA was used for subsequent steps. Ribosomal RNA was removed using a Ribo-Zero rRNA Removal Kit (Illumina, San Diego, CA, USA) according to the manufacturer’s instructions with a modification that included combining magnetic beads from the yeast and bacteria kit as follows: 0.5x Yeast Removal Solution, 0.25x Gram Negative Bacteria Removal solution, and 0.25x Gram Positive Bacteria Removal Solution. Strand-specific and barcode indexed RNA-seq libraries were then generated using the Kapa RNA-seq Hyper kit (Kapa Biosystems, Cape Town, South Africa) following the instructions of the manufacturer. The fragment size distribution of the libraries was verified via micro-capillary gel electrophoresis on a Bioanalyzer 2100. The libraries were quantified by fluorometry on a Qubit fluorometer and pooled in equimolar ratios. The pool was quantified by qPCR with a Kapa Library Quant kit (Kapa Biosystems) and sequenced on 1 lane of an Illumina HiSeq 4000 (Illumina, San Diego, CA, USA) with single-end 100 bp reads. The sequencing was carried out at the DNA Technologies and Expression Analysis Cores at the UC Davis Genome Center.

Resulting sequences from metatranscriptomic analysis were annotated with the Metagenomics Rapid Annotation using Subsystems Technology (MG-RAST) server version 4.0.3^44^. Functional classification was performed using the SEED Subsystems database and taxonomic annotations up to genus level were performed using the RefSeq database with a maximum e-value cut-off of 10^−5^, minimum identity cut-off of 60% and minimum length of sequence alignment of 15 nucleotides. Abundance tables derived from MG-RAST (Data S3) were imported into R for downstream analyses. Some samples were excluded from further analyses as we either could not extract good quality RNA (extraction from shrub litter was difficult) or the quality of sequences obtained was poor (due to poor quality RNA or bad sequencing runs). Number of replicates (n) for each treatment combination was- grass ambient control: 4, grass ambient wet-up: 3, grass ambient dry-down: 4, grass reduced control: 4, grass reduced wet-up: 4, grass reduced dry-down: 4, shrub ambient control: 4, shrub ambient wet-up: 2, shrub ambient dry-down: 3, shrub reduced control: 4, shrub reduced wet-up: 2, and shrub reduced dry-down: 3.

### Metabolite extraction and analysis

Leaf litter samples (1 g) were placed in 50 mL tubes with the addition of LC-MS grade water (20 mL, Honeywell Burdick & Jackson, Morristown, NJ, USA). Two extraction controls were included at this step by adding 20 mL of water to empty tubes. All samples were extracted for 1 h on an orbital shaker (Orbital-Genie, Scientific Industries, Bohemia, NY, USA) at 200 rpm at 4°C followed by centrifugation for 15 min at 3220 x g at 4°C. Supernatants (8 mL) were filtered through 0.45 um syringe filters (Pall Acrodisc Supor membranes) into 15 mL tubes, frozen then lyophilized. Dried extracts were resuspended in 1 mL LC-MS grade methanol (Honeywell Burdick & Jackson, Morristown, NJ, USA) on ice, vortexed for 10 sec, sonicated for 10 min in an ice bath and placed at 4°C overnight. Samples were centrifuged for 15 min at 3220 x g at 10°C and further cooled at −20°C for 10 min. Supernatants (850 μL) were transferred to 1.5 mL Eppendorf tubes and dried down with a Savant SpeedVac SPD111V (Thermo Scientific, Waltham, WA, USA) for 1 h with a final resuspension in ice-cold methanol containing internal standards (200 μL) and filtered through 0.22 μm centrifugal membranes (Nanosep MF, Pall Corporation, Port Washington, NY, USA) by centrifuging at 10,000 x g for 5 min. Samples (50 μL) were transferred to LC-MS vials for metabolomics analysis.

Extracts were analyzed using normal-phase LC-MS using a HILIC-Z column (150 mm × 2.1 mm, 2.7 μm, 120Å, Agilent Technologies, Santa Clara, CA, USA) on an Agilent 1290 series UHPLC. The two mobile phases for metabolite separation were 5 mM ammonium acetate in 0.2% acetic acid (A) and 95% acetonitrile with 5 mM ammonium acetate in 0.2% acetic acid (B) at a flow rate of 0.45 mL/min with the following gradient: 100% B for 1 min, decreased to 89% by 11 min, down to 70% by 15.75 min and 20% by 16.25 min, held until 18.5 min then back to 100% B by 18.6 min for a total runtime of 22.5 min. The column temperature was maintained at 40°C. MS data were collected on a Thermo Q Exactive (Thermo Fisher Scientific, Waltham, MA, USA) and MS-MS data were collected using collision energies of 10-40 eV. Metabolomics data were analyzed using Metabolite Atlas^45^ with in-house Python scripts to obtain extracted ion chromatograms and peak heights for each metabolite (Data S4). Metabolite identifications were verified with authentic chemical standards and validated based on three metrics (accurate mass less than 15 ppm, retention time within 1 min, and MS/MS fragment matching). Data from internal standards and quality control samples (included throughout the run) were analyzed to ensure consistent peak heights and retention times. One grass reduced control sample was lost during the extraction process, otherwise the number of replicates (n) for each treatment combination was 4.

### Ordination analysis and diversity indices

Rarefaction, ordination and diversity analyses of metatranscriptomics-derived functions and taxonomic units as well as metabolites were performed using the vegan package^46^, and NMDS plots were generated using the ggplot2 package^47^ under the R software environment 3.4.2^48^. Treatment effects were estimated using permutational multivariate analysis of variance (PERMANOVA) based on Bray Curtis dissimilarity distance between samples with the R-vegan function adonis. Shannon diversity index was calculated with rarefied functions and taxonomic units. Two-way factorial ANOVA and post hoc Tukey honest significant differences (HSD) tests were performed to ascertain the effect of treatments on functional and taxonomic α diversity.

### Gene expression analysis

Changes in gene expression on pulse wetting were normalised and quantified using differential expression analysis of RNA-Seq data and visualised using volcano plots, both implemented with DESeq2 R package^49^. The pairwise analysis was performed separately for grass ambient, grass reduced, shrub ambient, and shrub reduced precipitation treatments using control and wet-up samples. Genes were considered to be significantly up or downregulated in response to wetting if the p value of the differential analysis was smaller than 0.05 and fold change was more than double or half, respectively.

### Treatment effect size and indicator analysis

Since the effect of short-term wet-up and dry-down on gene expression and metabolite production was smaller in comparison to the differences across vegetation and precipitation treatments, we used samples from all short-term treatments to measure the impact of vegetation type and long-term field precipitation manipulation on community physiology. The reduced precipitation effect on the abundances of metatranscriptomics-derived functions at the upper level of SEED Subsystems classification was measured as the ratio of sum-normalised transcript abundance in level 1 classes in reduced and ambient precipitation treatments. One-way ANOVA was used to ascertain if this effect was significant.

To identify those transcripts that were significantly enriched in either reduced or ambient precipitation treatments in the two litter types, we used pairwise indicator species analysis as implemented within the R library labdsv^50^. The indval score for each transcript is the product of the relative frequency and relative average abundance within each treatment, and significance (p) was calculated through random reassignment of groups (1000 permutations). This indval score ranges from 0 to 1; maximum score of 1 for a transcript denotes that it is observed in all samples of only one treatment group. Transcripts above a threshold indval score of 0.65 and with a p value smaller than 0.05 were considered significant (Data S1). Indval score was chosen to have enough stringency while also aiming to obtain a manageable and meaningful number of functional indicators. To identify vegetation specific transcript indicators, we performed the indicator analysis on transcripts from ambient grass and ambient shrub treatments. We observed thousands of grass and shrub specific indicators. The threshold indval score for significance was raised to 0.7 for grass-shrub pairwise comparison to reduce the number of vegetation-specific indicators (Data S2). The frequency of indicator transcripts in level 1 functional classes was obtained using R plyr package^51^.

### Heatmap visualisations

Metabolite abundances were visualised using the heatmap function in R. Metabolites that were significantly higher in either ambient or reduced precipitation treatments across both litter types were identified using pairwise indicator species analysis as described above. Here, we did not separate samples from grass and shrub litter in order to reduce picking of litter-specific plant-derived metabolites as indicators. Metabolites that were significantly higher in wet-up treatment relative to the control were also identified using the same analysis. Metabolites above a threshold indval score of 0.6 and with a p value smaller than 0.05 were considered significant. A lower indval score was used here as higher stringency gave very few indicators.

### Correlation analysis and visualisation

Correlations between metabolite abundances were performed to quantify the identified tradeoffs between traits related to growth and stress tolerance. We tested for negative bivariate correlations between significantly enriched metabolites in either reduced or ambient precipitation treatments across both litter types. Correlation matrix visualization was done using ggcorrplot package in R^52^, separating samples from grass and shrub litter. Pearson correlation coefficient was used as a measure of the linear dependence between metabolites. From this correlation analysis, we chose key relevant metabolites that represent the clusters of significantly enriched metabolites to further visualize the metabolic trade-offs across and between treatments. These key metabolites were aspartic acid and adenosine (growth indicators), and ectoine and 5-oxo-proline (stress indicators). Psych Package in R^53^ was used to obtain bivariate scatter plots with linear model fits, correlation coefficients (r) and p values denoting the strength of relationships among the representative metabolites, separately for samples from grass and shrub litter.

## Acknowledgements

We acknowledge funding from the US Department of Energy (DOE) Genomic Science Program, BER, Office of Science project DE-SC0016410. Part of this work was performed at Lawrence Berkeley National Laboratory, under DOE contract No. DE-AC02-05CH11231. RNA sequencing was carried out at the DNA Technologies and Expression Analysis Cores at the UC Davis Genome Center, supported by NIH Shared Instrumentation Grant 1S10OD010786-01.

## Author contributions

AAM, JBHM, ELB and SDA designed research; AAM, CW, EM and SDA were involved in litter sampling, experimental setup and sample processing; AAM performed the RNA extractions; TS performed metabolite extraction and analysis; AAM performed bioinformatic and statistical analyses; JBHM, ELB and TN contributed reagents and analytical tools; AAM drafted the manuscript and all authors were involved in critical revision and approval of the final version.

## References

1. Dai, A. Increasing drought under global warming in observations and models. Nat. Clim. Chang. 3, 52–58 (2012).

2. Sherwood, S. & Fu, Q. A Drier Future? Science (80-.). 343, 737 LP–739 (2014).

3. Cook, B. I., Ault, T. R. & Smerdon, J. E. Unprecedented 21st century drought risk in the American Southwest and Central Plains. Sci. Adv. 1, e1400082 (2015).

4. Manzoni, S., Schimel, J. P. & Porporato, A. Responses of soil microbial communities to water stress: Results from a meta-analysis. Ecology 93, 930–938 (2012).

5. Schimel, J., Balser, T. C. & Wallenstein, M. Microbial stress-response physiology and its implications for ecosystem function. Ecology 88, 1386–1394 (2007).

6. Schimel, J. P. Life in Dry Soils: Effects of Drought on Soil Microbial Communities and Processes. Annu. Rev. Ecol. Evol. Syst. 49, 409–432 (2018).

7. Allison, S. D. et al. Microbial abundance and composition influence litter decomposition response to environmental change. Ecology 94, 714–725 (2013).

8. Hueso, S., García, C. & Hernández, T. Severe drought conditions modify the microbial community structure, size and activity in amended and unamended soils. Soil Biol. Biochem. 50, 167–173 (2012).

9. Lebre, P. H., De Maayer, P. & Cowan, D. A. Xerotolerant bacteria: surviving through a dry spell. Nat. Rev. Microbiol. (2017). doi:10.1038/nrmicro.2017.16

10. Bouskill, N. J. et al. Belowground response to drought in a tropical forest soil. I. Changes in microbial functional potential and metabolism. Front. Microbiol. 7, 1–11 (2016).

11. Placella, S. A., Brodie, E. L. & Firestone, M.K. Rainfall-induced carbon dioxide pulses result from sequential resuscitation of phylogenetically clustered microbial groups. Proc. Natl. Acad. Sci. 109, 10931 LP–10936 (2012).

12. Evans, S. E. & Wallenstein, M. D. Climate change alters ecological strategies of soil bacteria. Ecol. Lett. 17, 155–164 (2014).

13. Saetre, P. & Stark, J. M. Microbial dynamics and carbon and nitrogen cycling following re-wetting of soils beneath two semi-arid plant species. Oecologia 142, 247–260 (2005).

14. Blazewicz, S. J. et al. Growth and death of bacteria and fungi underlie rainfall-induced carbon dioxide pulses from seasonally dried soil Published by : Wiley on behalf of the Ecological Society of America Stable URL : http://www.jstor.org/stable/43494786 Growth and death of bac. Ecology 95, 1162–1172 (2014).

15. Barnard, R. L., Osborne, C. A. & Firestone, K. Responses of soil bacterial and fungal communities to extreme desiccation and rewetting. ISME J. 7, 2229–2241 (2013).

16. Kim, D. G., Vargas, R., Bond-Lamberty, B. & Turetsky, M. R. Effects of soil rewetting and thawing on soil gas fluxes: A review of current literature and suggestions for future research. Biogeosciences 9, 2459–2483 (2012).

17. Swenson, T. L., Karaoz, U., Swenson, J. M., Bowen, B. P. & Northen, T. R. Linking soil biology and chemistry in biological soil crust using isolate exometabolomics. Nat. Commun. 9, (2018).

18. Steven, B., Belnap, J. & Kuske, C. R. Chronic physical disturbance substantially alters the response of biological soil crusts to a wetting pulse, as characterized by metatranscriptomic sequencing. Front. Microbiol. 9, 1–17 (2018).

19. Kimball, S., Goulden, M. L., Suding, K. N. & Parker, S. Altered water and nitrogen input shifts succession in a southern California coastal sage community. Ecol. Appl. 24, 1390–1404 (2014).

20. Strickland, M. S., Osburn, E., Lauber, C., Fierer, N. & Bradford, M. a. Litter quality is in the eye of the beholder: initial decomposition rates as a function of inoculum characteristics. Funct. Ecol. 23, 627–636 (2009).

21. Martiny, J. B. H. et al. Microbial legacies alter decomposition in response to simulated global change. ISME J. 11, 490–499 (2017).

22. Malik, A. A. et al. Defining trait-based microbial strategies with consequences for soil carbon cycling under climate change. bioRxiv (2018).

23. Fierer, N., Barberán, A. & Laughlin, D. C. Seeing the forest for the genes: Using metagenomics to infer the aggregated traits of microbial communities. Front. Microbiol. 5, 1–6 (2014).

24. Malik, A. A., Thomson, B. C., Whiteley, A. S., Bailey, M. & Griffiths, R. I. Bacterial physiological adaptations to contrasting edaphic conditions identified using landscape scale metagenomics. MBio 8, (2017).

25. Warren, C. R. Response of osmolytes in soil to drying and rewetting. Soil Biol. Biochem. 70, 22–32 (2014).

26. Varin, T., Lovejoy, C., Jungblut, A. D., Vincent, W. F. & Corbeil, J. Metagenomic analysis of stress genes in microbial mat communities from Antarctica and the high Arctic. Appl. Environ. Microbiol. 78, 549–559 (2012).

27. Pardo, J. M., Cubero, B., Leidi, E. O. & Quintero, F. J. Alkali cation exchangers: roles in cellular homeostasis and stress tolerance. J. Exp. Bot. 57, 1181–1199 (2006).

28. Kakumanu, M. L., Cantrell, C. L. & Williams, M. A. Microbial community response to varying magnitudes of desiccation in soil: A test of the osmolyte accumulation hypothesis. Soil Biol. Biochem. 57, 644–653 (2013).

29. Tanaka, K. The proteasome: overview of structure and functions. Proc. Jpn. Acad. Ser. B. Phys. Biol. Sci. 85, 12–36 (2009).

30. Finn, S., Condell, O., McClure, P., Amézquita, A. & Fanning, S. Mechanisms of survival, responses, and sources of salmonella in low-moisture environments. Front. Microbiol. 4, 1–15 (2013).

31. Wood, J. M. Bacterial responses to osmotic challenges. J. Gen. Physiol. 145, 381 LP–388 (2015).

32. Halverson, L. J., Jones, T. M. & Firestone, K. Release of Intracellular Solutes by Four Soil Bacteria Exposed to Dilution Stress. Soil Sci. Soc. Am. J. 64, 1630–1637 (2000).

33. Kocharunchitt, C., King, T., Gobius, K., Bowman, J. P. & Ross, T. Integrated Transcriptomic and Proteomic Analysis of the Physiological Response of *Escherichia coli* O157:H7 Sakai to Steady-state Conditions of Cold and Water Activity Stress. Mol. Cell. Proteomics 11, M111.009019 (2012).

34. Malik, A. A. et al. Land use driven change in soil pH affects microbial carbon cycling processes. Nat. Commun. 9, 3591 (2018).

35. Berlemont, R. et al. Cellulolytic potential under environmental changes in microbial communities from grassland litter. Front. Microbiol. 5, 639 (2014).

36. Evans, S. E. & Wallenstein, M. D. Soil microbial community response to drying and rewetting stress: Does historical precipitation regime matter? Biogeochemistry 109, 101–116 (2012).

37. Allison, S. D. & Goulden, M. L. Consequences of drought tolerance traits for microbial decomposition in the DEMENT model. Soil Biol. Biochem. 107, 104–113 (2017).

38. Preece, C., Verbruggen, E., Liu, L., Weedon, J. T. & Peñuelas, J. Effects of past and current drought on the composition and diversity of soil microbial communities. Soil Biol. Biochem. 131, 28–39 (2018).

39. Matulich, K. L. & Martiny, J. B. H. Microbial composition alters the response of litter decomposition to environmental change. Ecology 96, 154–163 (2015).

40. Flemming, H.-C. et al. Biofilms: an emergent form of bacterial life. Nat. Rev. Microbiol. 14, 563 (2016).

41. Tiemann, L. K. & Billings, S. A. Changes in variability of soil moisture alter microbial community C and N resource use. Soil Biol. Biochem. 43, 1837–1847 (2011).

42. Killham, K. & Firestone, M. K. Proline transport increases growth efficiency in salt-stressed Streptomyces griseus. Appl. Environ. Microbiol. 48, 239–241 (1984).

43. Esch, E. H., King, J. Y. & Cleland, E. E. Foliar litter chemistry mediates susceptibility to UV degradation in two dominant species from a semi-arid ecosystem. (2019).

44. Meyer, F. et al. The metagenomics RAST server - a public resource for the automatic phylogenetic and functional analysis of metagenomes. BMC Bioinformatics 9, 386 (2008).

45. Yao, Y. et al. Analysis of metabolomics datasets with high-performance computing and metabolite atlases. Metabolites 5, 431–442 (2015).

46. Oksanen, J. et al. vegan: Community Ecology Package. R package version 2.3-0. (2015).

47. Wickham, H. ggplot2: Elegant Graphics for Data Analysis. (2016).

48. R Development Core Team. R: A language and environment for statistical computing. (2017).

49. Love, M. I., Huber, W. & Anders, S. Moderated estimation of fold change and dispersion for RNA-seq data with DESeq2. Genome Biol. 15, 550 (2014).

50. Roberts, D. W. labdsv: Ordination and Multivariate Analysis for Ecology. (2016).

51. Wickham, H. The Split-Apply-Combine Strategy for Data Analysis. J. Stat. Software; Vol 1, Issue 1 (2011). doi:10.18637/jss.v040.i01

52. Kassambara, A. Visualization of a Correlation Matrix using ‘ggplot2’. (2018).

53. Revelle, W. Procedures for Psychological, Psychometric, and Personality Research. (2019).

