## Supplementary Information for "Physiological adaptations of leaf litter microbial communities to long-term drought"

Malik et al.

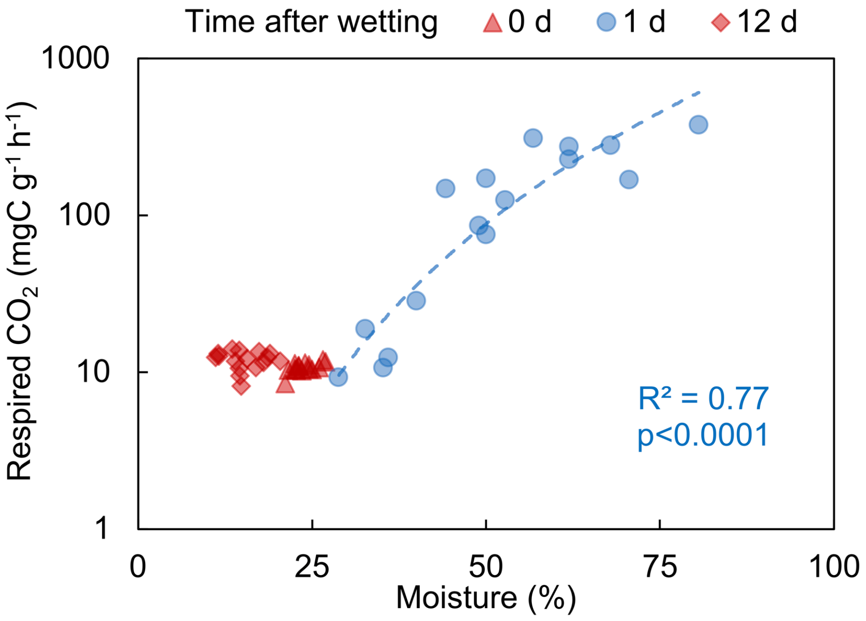


Figure S1: Rate of respired $CO_2$ across the wet-up (day 1) and dry-down (day 12) treatments relative to control (day 0). Best fitting regression line for the moisture-respired $CO_2$ rate relationship: $y = 1E\text{-}05x^{4.04}$.

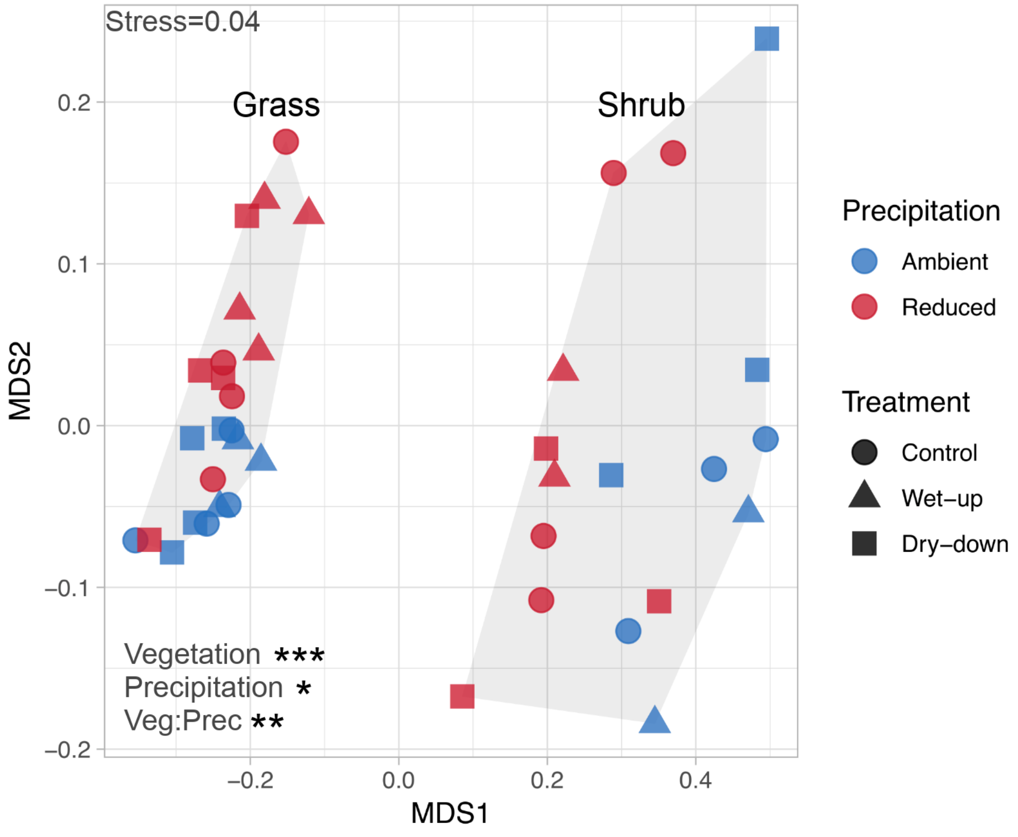


Figure S2: Two dimensional NMDS ordination of communities identified at the genus level from metatranscriptomics based on taxonomic annotations of functional genes. Asterisks mark the significance of treatments that cause clustering of similar samples based on Bray-Curtis dissimilarity index analysed using permutational multivariate analysis of variance (PERMANOVA); *** $p < 0.001$, ** $p < 0.01$, * $p < 0.05$.

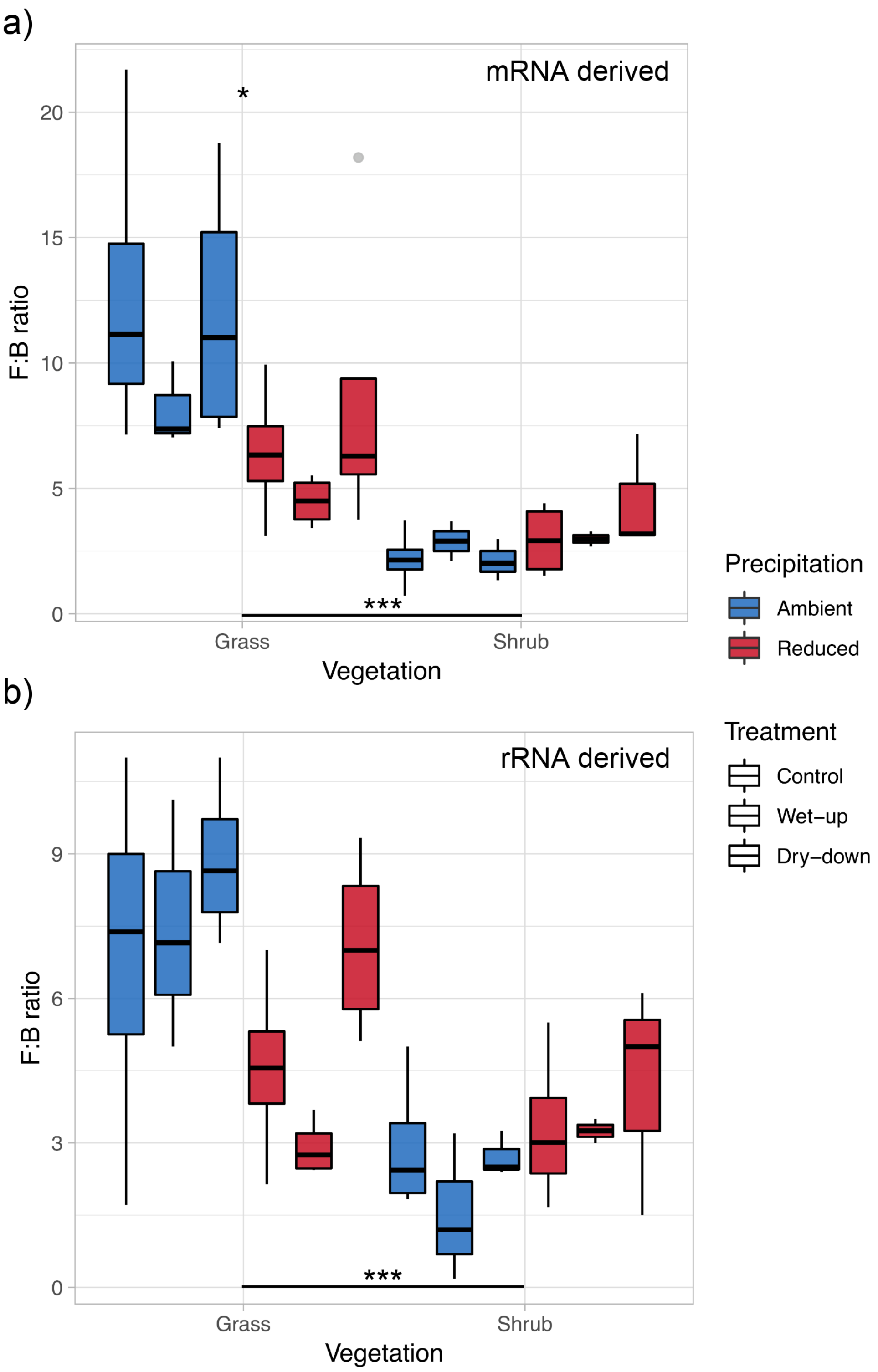


Figure S3: Community fungal:bacterial (F:B) ratio estimated as (a) the ratio of mRNA sequences assigned to fungi and bacteria (b) the ratio of 18S and 16S rRNA abundances in total RNA extracts; n=3-4. Most of the mRNA (~ 97%) was annotated to fungi or bacteria, therefore we assumed that majority of 16S belonged to bacteria and 18S to fungi. Asterisks mark the significance of differences between groups as analysed by Tukey's multiple comparison test (*** $p < 0.001$, ** $p < 0.01$, * $p < 0.05$).

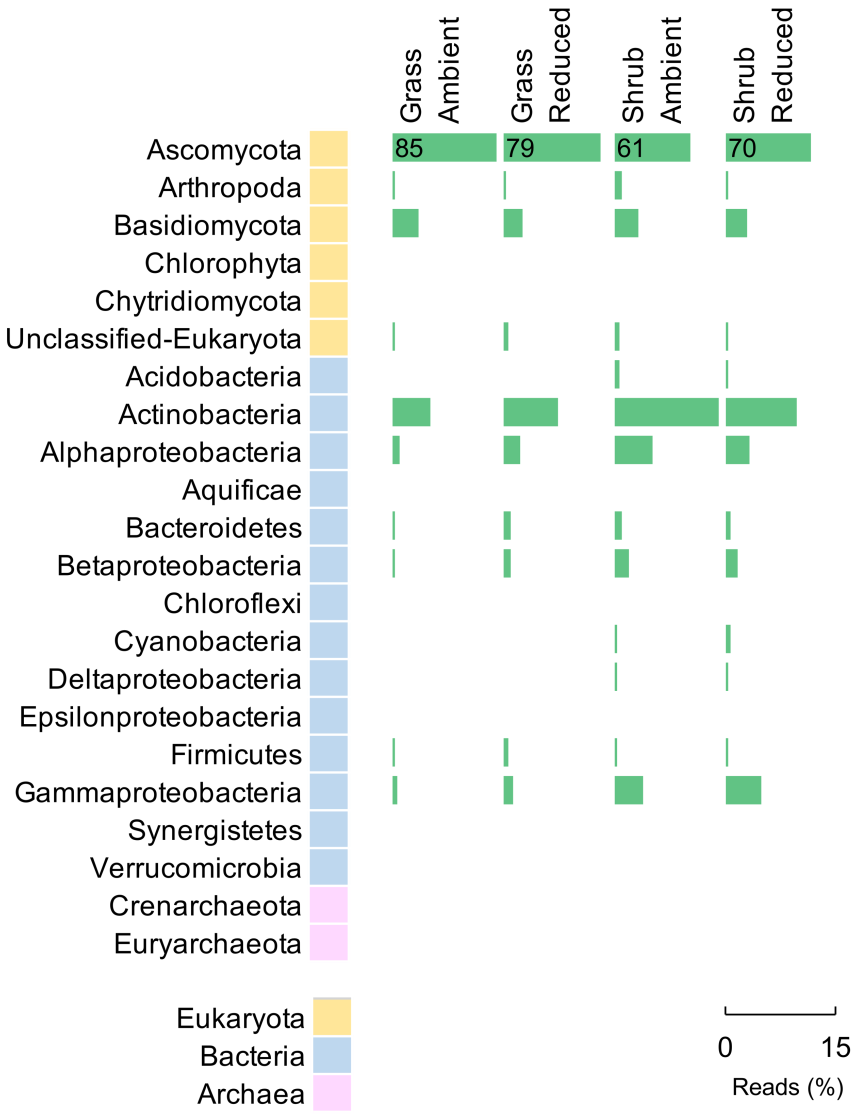


Figure S4: The mean proportion of various phyla/sub-phyla in communities from the litter types and precipitation treatments. Ascomycota was the most abundant phylum, it was disproportionately higher than the rest of the phyla and therefore has been plotted on a separate scale with its abundance labelled on the bar.

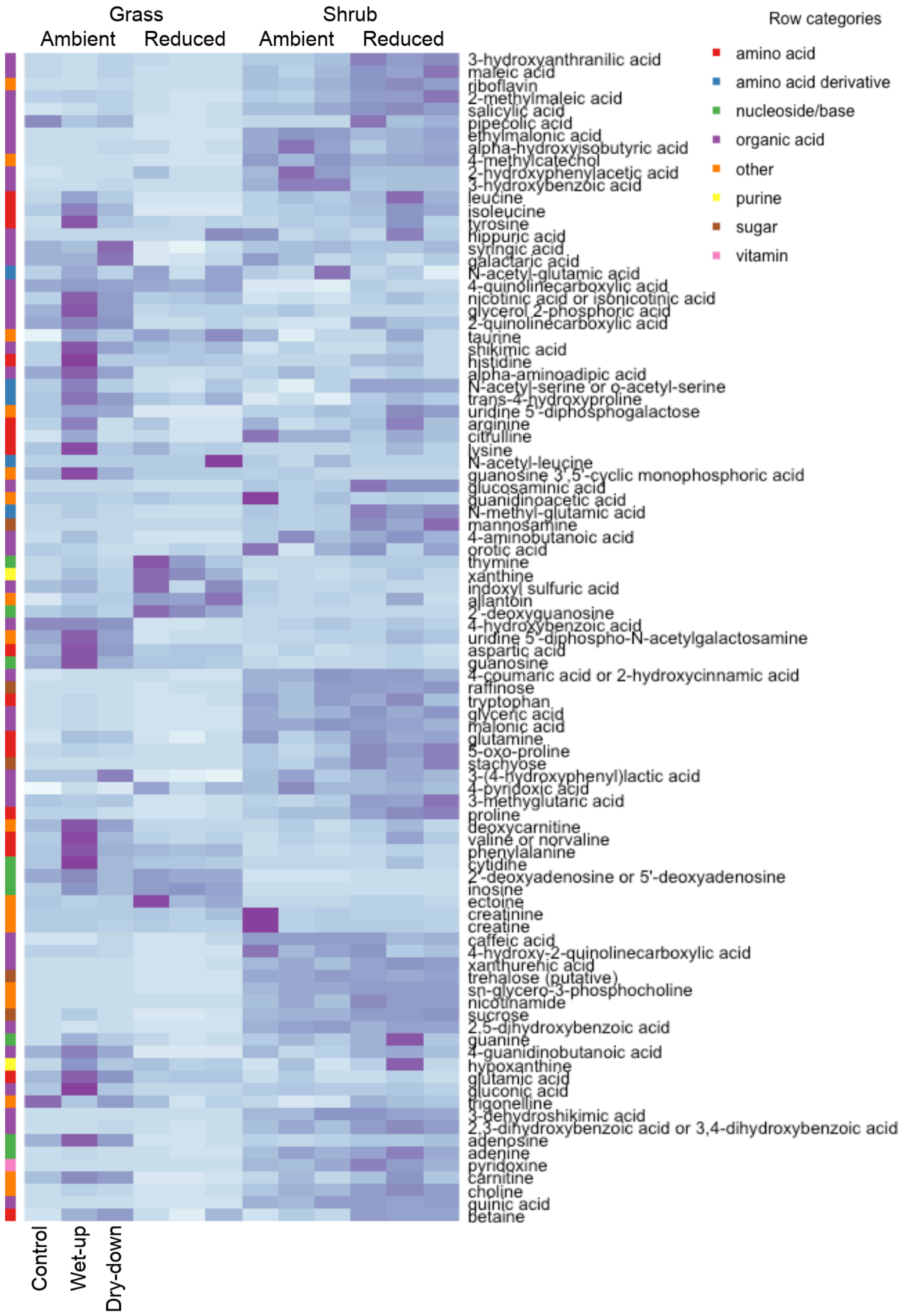


Figure S5: Heatmap of mean peak heights (n=3-4) of all identified metabolites across treatments.

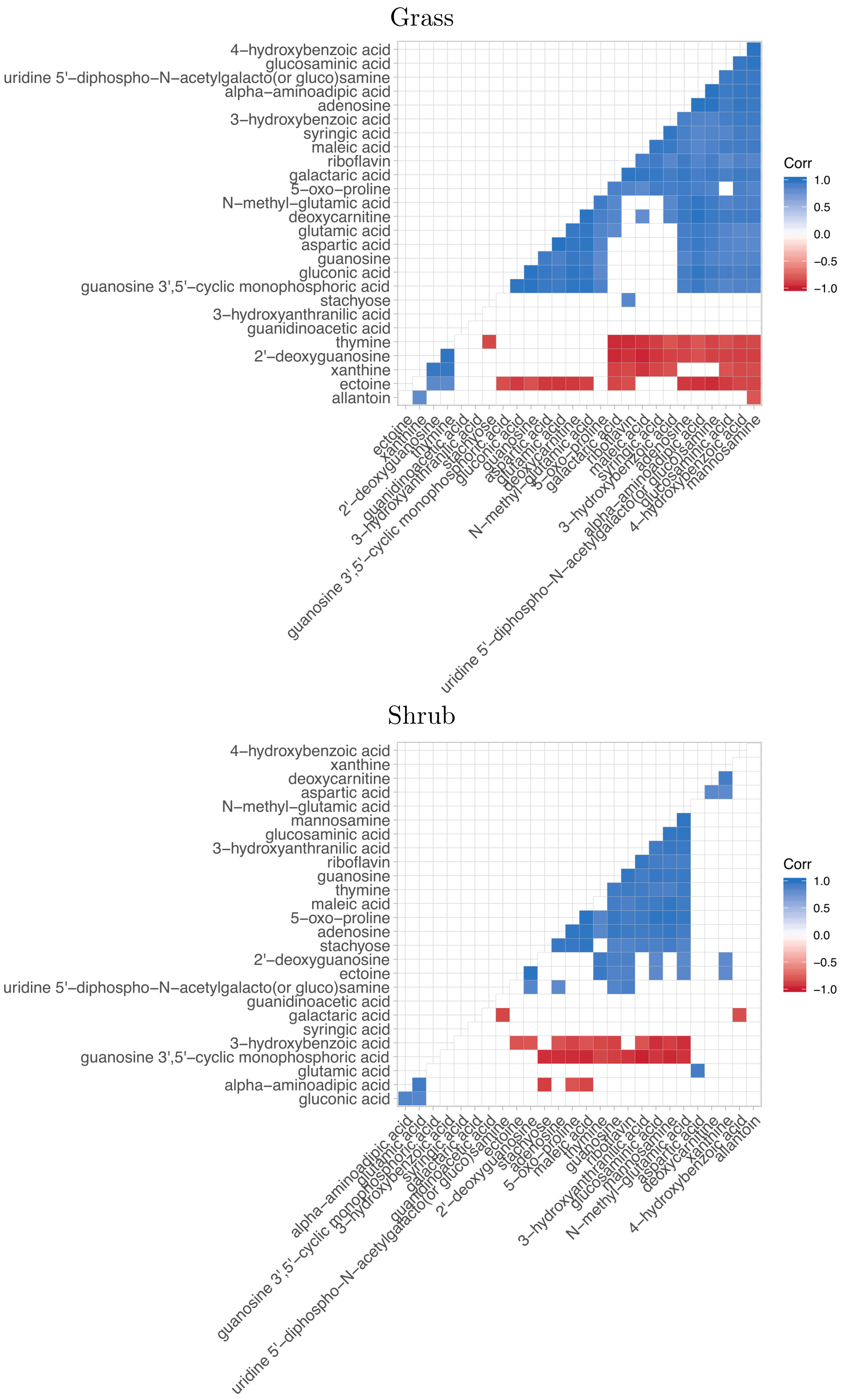


Figure S6: Correlations between metabolites significantly enriched in either ambient or reduced precipitation treatments across the two vegetation types. Empty boxes signify that the correlations are not significant.

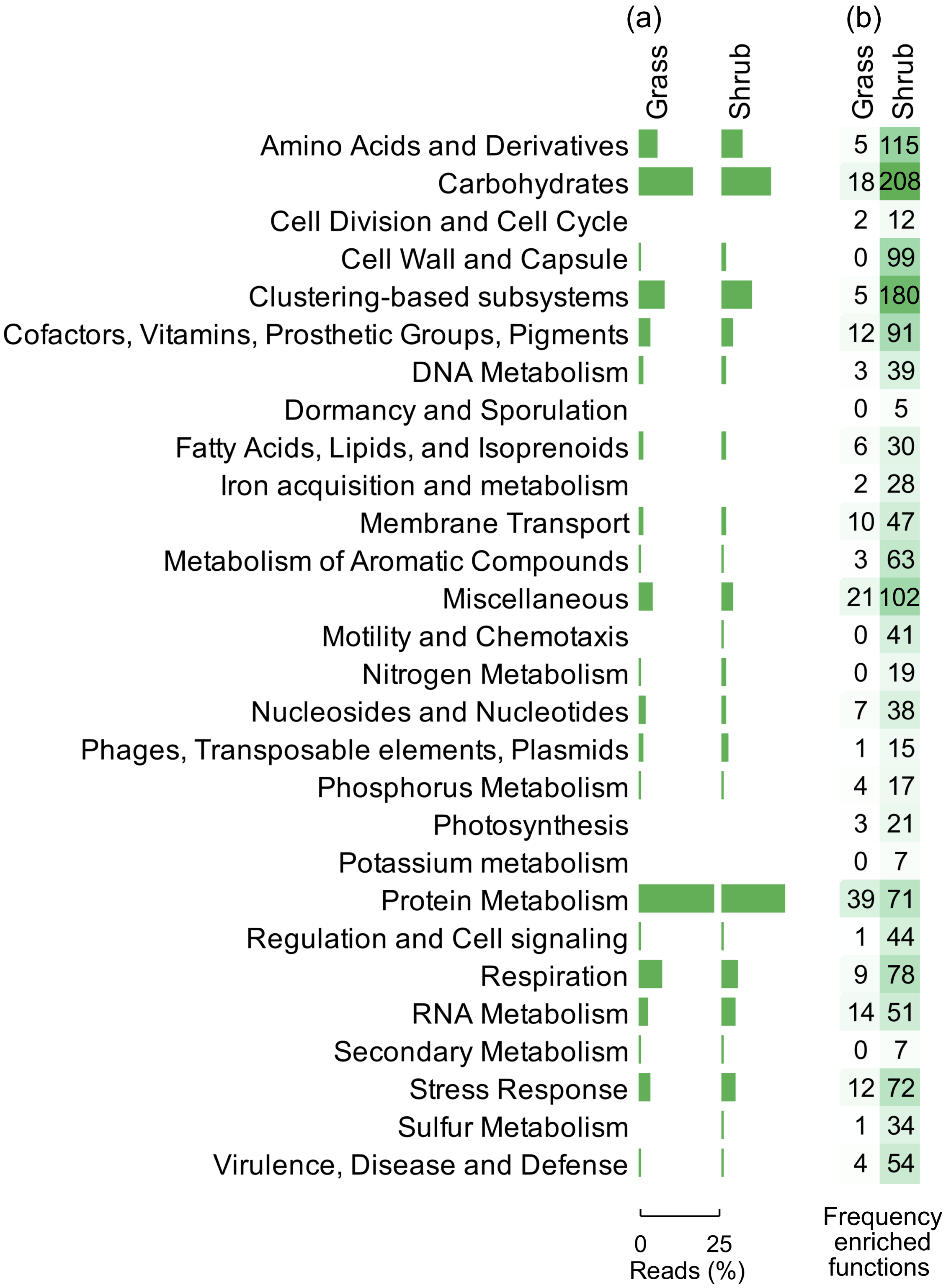


Figure S7: (a) Mean relative abundance of transcripts at level 1 of Subsystems classification in grass ambient and shrub ambient precipitation treatments. (b) Frequency of significant transcript indicators in the upper level functional groups (level 1 class) across vegetation types with ambient precipitation. These indicators were unique to or enriched in communities in either grass ambient or shrub ambient precipitation treatments and represent functional indicators of grass or shrub ecosystems.

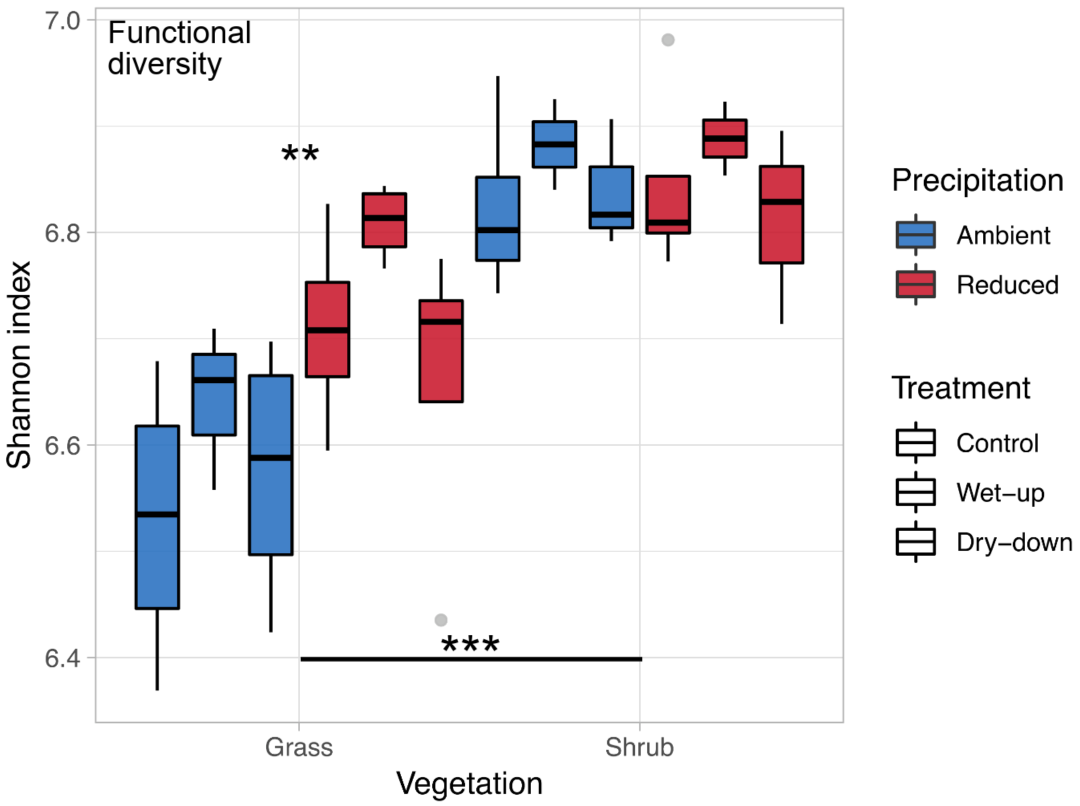


Figure S8: Functional α-diversity derived from metatranscriptomics across vegetation types, precipitation and pulse wet-up/dry-down treatments; n=3-4. Asterisks mark the significance of differences between groups as analysed by Tukey's multiple comparison test (*** $p < 0.001$, ** $p < 0.01$, * $p < 0.05$).

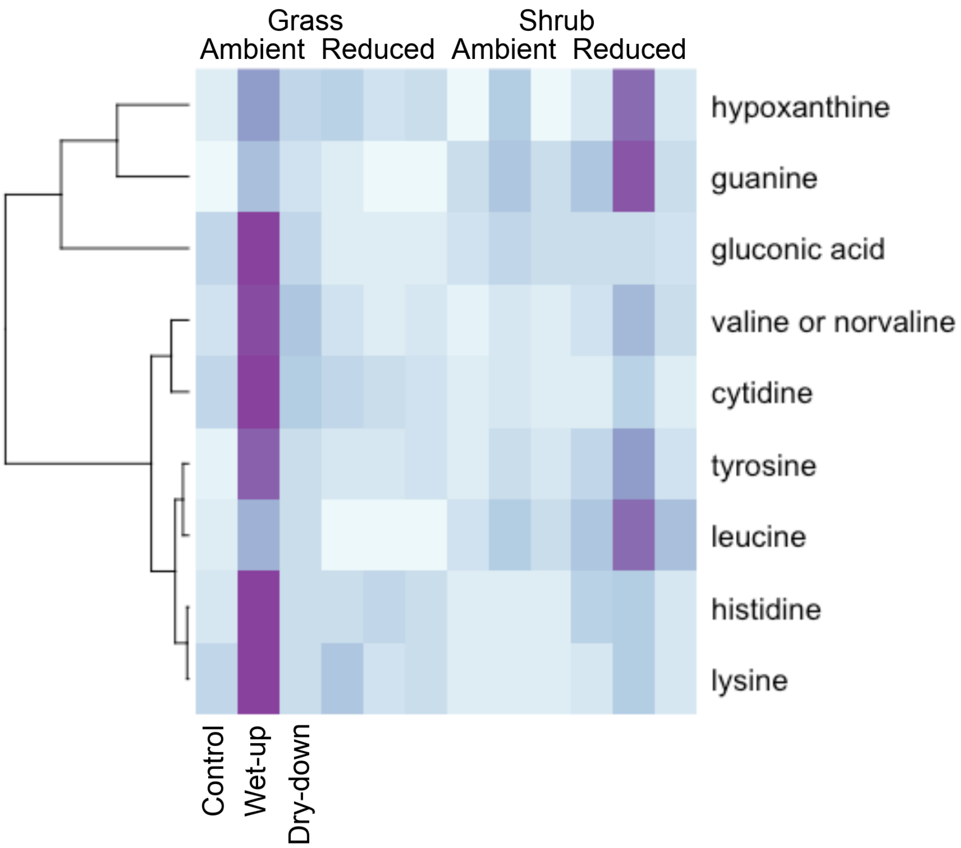


Figure S9: Heatmap of mean peak heights (n=3-4) which relates to the abundance of metabolites that were significantly higher ($p<0.05$) in the wet-up treatment.
